# *De Novo* Variants in *MRTFB* have gain of function activity in *Drosophila* and are associated with a novel neurodevelopmental phenotype with dysmorphic features

**DOI:** 10.1101/2022.12.06.518921

**Authors:** Jonathan C. Andrews, Jung-Wan Mok, Oguz Kanca, Sharayu Jangam, Cynthia Tifft, Ellen F. Macnamara, Bianca Russell, Lee-kai Wang, Undiagnosed Diseases Network, Stanley F. Nelson, Hugo J. Bellen, Shinya Yamamoto, May Malicdan, Michael F. Wangler

## Abstract

Myocardin-Related Transcription Factor B (*MRTFB*) is an important transcriptional regulator which promotes the activity of an estimated 300 genes during different stages of development. Here we report two pediatric probands with *de novo* variants in *MRTFB* (R104G and A91P) and mild dysmorphic features, intellectual disability, global developmental delays, speech apraxia, and impulse control issues. As the *MRTFB* protein is highly conserved between vertebrate and invertebrate model organisms, we generated a humanized *Drosophila* model expressing the human *MRTFB* protein in the same spatial and temporal pattern as the fly gene. Expression of the human *MRTFB^R104G^* variant using a *mrtf-T2A-GAL4* line proved to be embryonic lethal. Additional phenotypes were also identified by expressing the *MRTFB^R104G^* and *MRTFB^A91P^* variant in a subset of *Drosophila* tissues. Notably, expression within wing tissues resulted in an expansion of intervein tissue, wing vein thickening, shortening or loss of wing veins, and blistering. The *MRTFB^R104G^* and *MRTFB^A91P^* variants also display a decreased level of actin binding within critical RPEL domains, resulting in increased transcriptional activity and changes in the organization of the Actin cytoskeleton. These changes were not observed in flies expressing two additional candidate variants, *MRTFB^N95S^*and *MRTFB^R109Q^*, highlighting that the location of the mutation within the 2nd RPEL domain is critical to the pathogenicity of the variant. These changes suggest that the *MRTFB^R104G^* and *MRTFB^A91P^* alleles we have identified affect the regulation of the protein and that these variants in *MRTFB* underly a novel neurodevelopmental disorder.

## Main Text

Myocardin-Related Transcription Factor B (*MRTFB*) is one of three members of the myocardin family^1–3^, a group of transcription factors known to contribute to proper cardiac development, smooth muscle differentiation, and the regulation of synaptic morphology^4–11^. The three members of this family, *MYOCD, MRTFA* and *MRTFB*, are located at different chromosomal loci but share several distinct structural features, including an N-terminal domain, a basic region, a B-box-like region, a glutamine-rich region, a SAP domain, a leucine zipper and a transcription activation domain^1,12–14^ (FIG. 1A). The N-terminal domain contains a series of three RPEL subdomains which are highly conserved between different species and are known to play an important role in the regulation of *MRTFB* activity^15^. Each of the three RPEL domains are capable of binding with G-actin, with the third RPEL domain showing the weakest affinity for actin binding^16,17^. X-ray crystallography studies of *MRTFA* have also revealed that the space between RPEL domains contributes to actin binding, and as *MRTFB* shares these residues, the spacer regions present in *MRTFB* are presumed to play a similar role^16,17^. The presence of bound actin in the RPEL domains and nearby spacer regions plays a critical role in determining the cellular localization of *MRTFA* and *MRTFB* by decreasing their binding affinity with importin α1/β1 heterodimers, and subsequently reducing the rate of nuclear import (Fig. 1B)^18^. Similarly, actin binding promotes the nuclear export of myocardin family members via exportin Crm1^17^. These two processes highlight why *MRTFB* signaling is particularly sensitive to the G-Actin concentration in cells and why the regulation of signaling pathways which influence actin polymerization, such as Rho signaling, can influence the subcellular location of *MRTFB*^19,20^.

**Figure 1.**
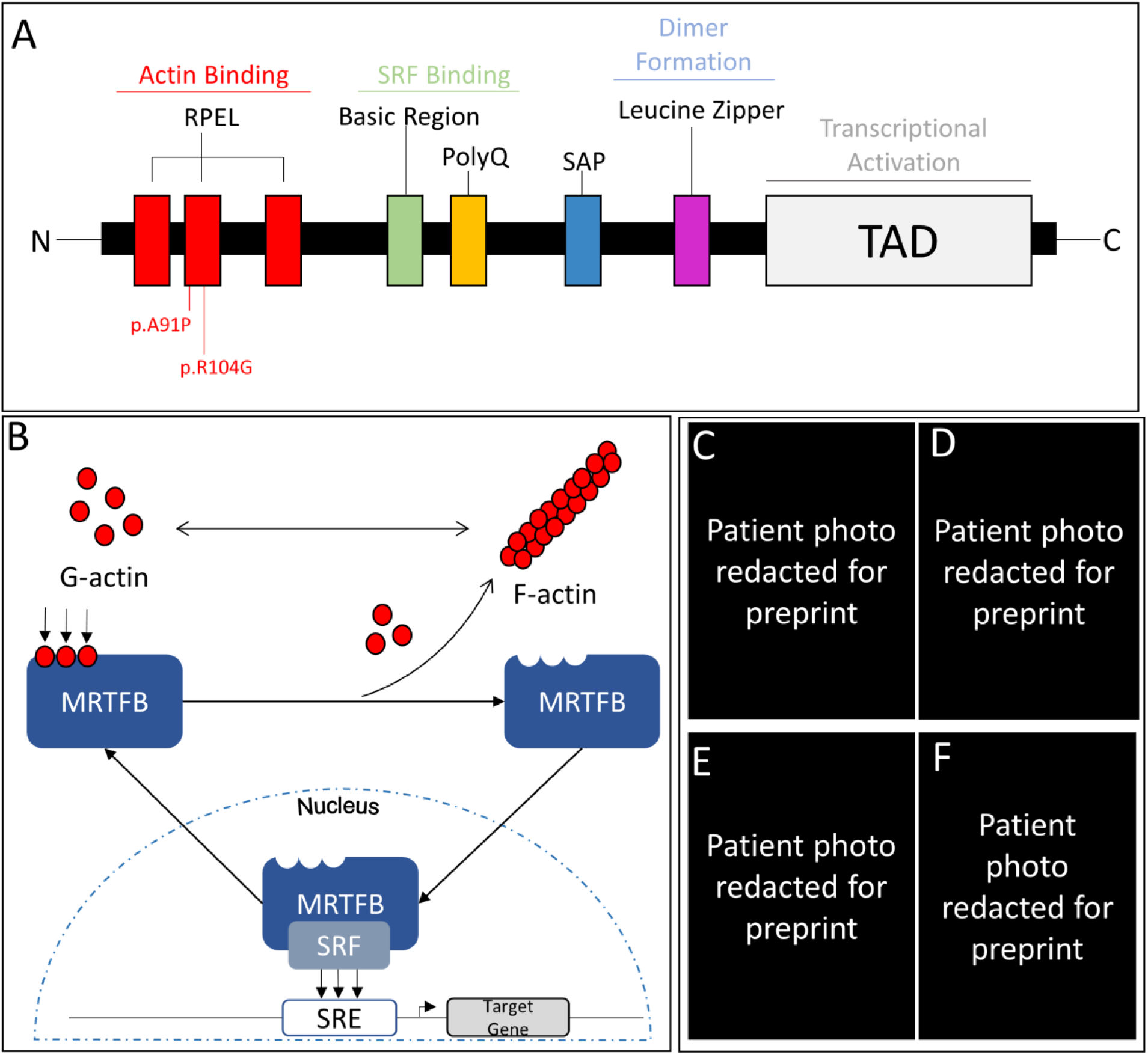
**(A)** Diagram of the structure of MRTFB. MRTFB contains a series of RPEL motifs, a basic region, a polyglutamine repeat (polyQ), a SAP domain, a leucine zipper domain, and a transcriptional activation domain. Both proband variants are found within the 2^nd^ RPEL domain. **(B)** While bound to G-actin, MRTFB remains within the cytoplasm. As cytoplasmic G-actin levels fall, MRTFB becomes unbound from actin, and is capable of translocating into the nucleus. Once within the nucleus, MRTFB forms a complex with its cofactor SRF, and can act as a translation factor for ~300 different genes. **(C-F)** Probands exhibit facial dysmorphology. The presence of synophrys and low set ears can be observed in both probands, while widely spaced teeth can be observed in proband 1. Proband 2 displays epicanthal folds, downslanting palpebral fissures, midface hypoplasia, a depressed nasal bridge, and tubular nose with bulbous tip.

The ability of *MRTFB* to act as a transcriptional regulator is primarily determined by its subcellular localization and its ability to bind its cofactor, *serum response factor (SRF)^17^. SRF*, in conjunction with its binding partners *MRTFA* or *MRTFB*, is responsible for the regulation of an estimated 300 different genes, including developmentally critical components of the actin cytoskeleton (*ACTB, ACTG1, GSN*) and synaptic activity (*CDK5R1, CDK5, RYR1, RYR3, CLTC, DLG4, ARC*)^21–24^. The activity of the *MRTFB-SRF* complex is particularly critical during early development, where mouse models have demonstrated its role in shaping development of the heart, lungs, liver, and brain^3–8,25^. Early global knock-out mouse models of *MRTFB* noted that the loss of *MRTFB* resulted in vascular malformations and subsequently embryonic lethality, which was not rescued by expression of other members of the myocardin family^6,26^. *Drosophila*, by comparison, possess a single gene, *Mrtf*, which is orthologous to the three vertebrate genes and has been implicated in multiple processes including cytoskeletal organization, cell migration, cell stretching, as well as learning and memory processes in an Actin-dependent manner^27–30^. Homozygous knock out models of the single *Drosophila* orthologue proved to be lethal during the 1^st^ instar stage due to defects in the growth and development of the tracheal system^28^. The possibility that *MRTFB* may cause disease has been suggested as a single case of three infants from the same family carrying transheterozygous alleles of *MRTFB* which were suggested, but not causally proven, to lead to early perinatal lethality and extreme microcephaly^31^.

Here we report two individuals with missense *de novo* variants in *MRTFB* who have overlapping phenotypes including intellectual disability, global developmental delay, dysmorphic features, and speech apraxia. Both probands were identified through the Undiagnosed Disease Network and were subsequently evaluated and sequenced under clinical protocols approved by the NIH and UCLA respectively. The individuals have overlapping phenotypes and neither have alternative genetic explanations from the diagnostic studies (Table 1). Notably the facial dysmorphology displayed significant overlap between the two probands, with widely spaced eyes, a prominent forehead, broad nasal bridge, widely spaced teeth, and synophrys (Table 2). Proband 1 (FIG. 1C–D), was diagnosed with motor and language delay at age 3, as well as an enlarged liver. Proband 2 (FIG. 1E–F) was noted to be developmentally delayed at 6 months, and was later diagnosed with ADHD and anxiety, in addition to language delay. For full details on both probands, see supplemental text 1. In the case of proband 1, clinical quartet genome sequencing, with that of an unaffected sibling, identified a *de novo* NM_014048.4; c.310C>G; p.R104G missense change in *MRTFB*. Proband 2 had trio sequencing which identified a *de novo* NM_014048.4; c.271G>C; p.A91P missense change in *MRTFB*. Neither the *MRTFB^R104G^* nor the *MRTFB^A91P^* variant was observed in gnomAD or ExAC databases and both were identified as damaging by multiple variant effect prediction tools, including CADD, SIFT, PolyPhen, and PROVEAN (Table 3)^32–35^.

**Table 1.** Summary of clinical features of probands with *de novo MRTFB* variants.

| Clinical Feature | Proband 1 | Proband 2 |
| --- | --- | --- |
| Gross Motor Delay (HP:0001270) | + | + |
| Fine Motor Delay (HP:0010862) | + | + |
| Language Delay (HP:0002474) | + | + |
| Hypotonia (HP:0001252) | + | + |
| Intellectual Disability, Severe (HP:0010864) | + | + |
| Enlarged Liver (HP:0002240) | + |  |
| Difficult Sleep (HP:0002360) | + |  |
| Apraxia (HP:0002186) | + |  |
| Speech Apraxia (HP:0011098) |  | + |
| Autism Spectrum Disorder (HP:0000717) |  | + |
| ADHD (HP:0007018) |  | + |
| Anxiety (HP:0000739) |  | + |
| Strabismus (HP:0000486) |  | + |

**Table 2.**
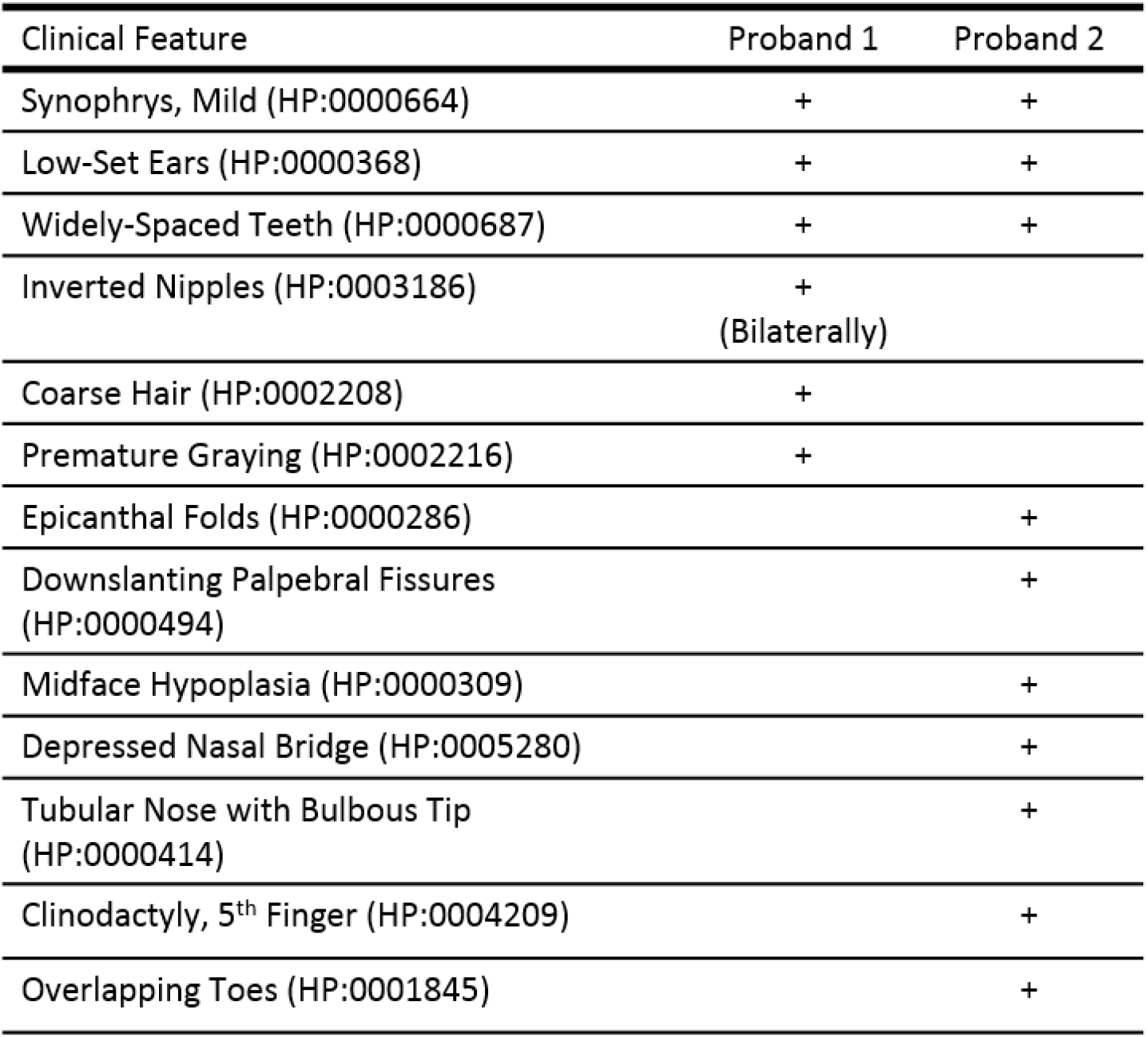
Dysmorphic features of probands with *de novo MRTFB* variants.

**Table 3.** *De Novo MRTFB* Variants

| Table 3. *De Novo MRTFB* Variants |  |  |
| --- | --- | --- |
| Variant Details | Proband 1 | Proband 2 |
| cDNA (NM_014048.4) | c.310C>G | c.271G>C |
| gDNA (Chr16,GRCh37) | g.14307435C>G | g. 14306261G>C |
| Protein | p.R104G | p.A91P |
| CADD Phred Score | 26.7 | 28.9 |
| SIFT | Damaging (Score = 0.02) | Damaging (Score = 0.00) |
| PolyPhen | Probably Damaging | Probably Damaging |
| PROVEAN | Deleterious (Score = -6.440) | Deleterious (Score = -4.244) |

To determine the functional consequences of the *MRTFB^R104G^* and *MRTFB^A91P^* variants, we utilized *Drosophila melanogaster* as a model system. An initial survey of databases was performed using MARRVEL (model organism aggregated resources for rare variant exploration) to assemble data from multiple online sources including ExAC, gnomAD, and OMIM^36–38^. In *Drosophila, MRTFB* has a single orthologous gene, *Mrtf*, with moderate homology (DIOPT score of 9/15, 24% amino acid identity and 37% similarity)^39^. Notably, the 2nd RPEL domain is highly conserved between humans and flies. We generated a new T2A-GAL4 allele for *Mrtf* by inserting the T2A-Gal4 sequence into the intron between the fifth and sixth exon of the endogenous locus of *Mrtf*^40^. This allele allows for sensitive detection of the *Mrtf* expression pattern and a “humanization” strategy where the human MRTFB protein is expressed in the same spatial and temporal pattern as the endogenous *Mrtf* (FIG 2A)^41^. Crosses which generate homozygous *mrtf-T2A-GAL4* flies were found to be lethal and *mrtf-T2A-GAL4/mrtf^ko^* animals carrying a null allele are also lethal, suggesting that the *mrtf-T2A-GAL4* line is a severe loss of function allele. Upon crossing the *Mrtf-T2A-GAL4* with a *UAS-nls:mCherry*, we observed that *Mrtf-T2A-GAL4* is widely expressed throughout both adult males and females (FIG 2B–C), in agreement with high-throughput expression profiling data^42^. As both of the probands also displayed a number of neurological symptoms, we dissected out the brains of adult *mrtf-T2A*; *UAS-nls:mCherry* animals and stained for the neuronal marker *elav* and the glial marker *repo* (FIG 2D–I). As shown, *Mrtf* is expressed in roughly 70% of the neurons of the adult brain and 20% of glia.

**Figure 2.**
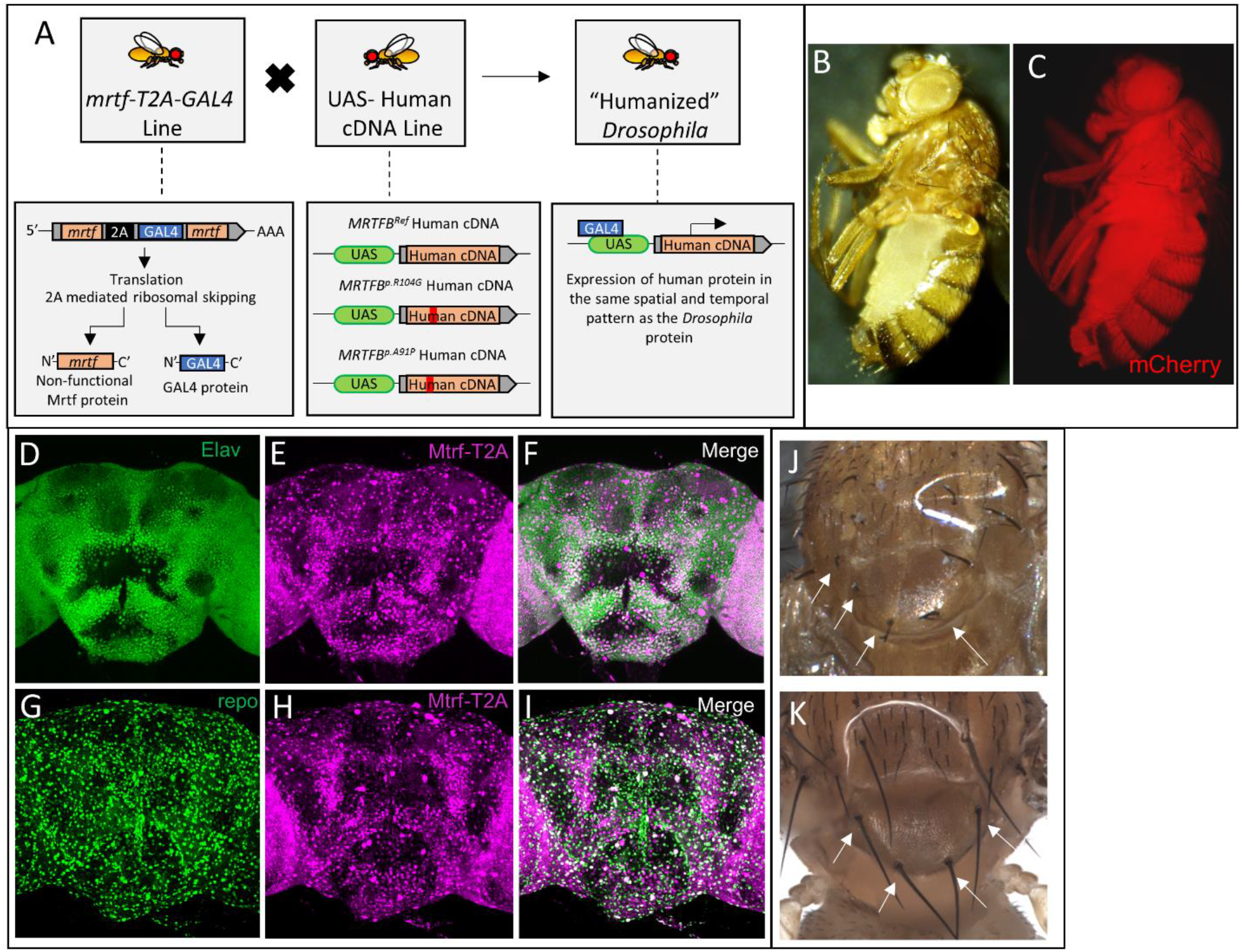
**(A)** Humanization strategy. A T2A-GAL4 line was generated to both disrupt gene function and express GAL4 in the same spatial and temporal pattern as the fly gene. By driving the expression of human reference and variant cDNA in the same pattern as the fly gene, we can attempt rescue with a “humanized” *Drosophila*. **(B)** UAS-GFP; *mrtf-T2A* fly under white-light illumination. **(C)** Expression of *UAS-nls-mCherry* under the control of *mrtf-T2A*. Expression is near-ubiquitous in the adult fly. **(D-I)** Expression of the *mrtf-T2A* line within the adult *Drosophila* brain. **(D)** Co-staining with Elav identifies neurons **(E)** *UAS-nls::mCherry* driven by *mrtf-T2A* **(F**) Merge of staining and RFP indicates that *mrtf* is expressed in ~70% of the neurons. **(G)** Co-staining with REPO identifies glia **(H)** *UAS-nls::mCherry* driven by *mrtf-T2A*. **(I)** Merge of staining and RFP indicates that *mrtf* is expressed in ~20% of glia. **(J)** *mal-dΔ7/mrtf-T2A* flies display a kinked bristle phenotype indicating that the T2A line is a loss of function. Arrows highlight kinked bristles. **(K)** *mal-dΔ7/+* flies display normal bristles. Arrows highlight normal bristles.

*Mrtf* was previously shown to be important for the proper development of actin-rich structures, such as the bristles present on the notum, and homozygous *mal-d^Δ7^* mutants missing the first exon of *Mrtf* possess a distinct kinked-bristle phenotype^27^. *mrtf-T2A-GAL4/mal-d^Δ7^* mutants also possess this bristle phenotype (FIG 2J–K). Introduction of the UAS-reference (*MRTFB^Ref^*) failed to rescue the bristle phenotype of the *mrtf-T2A-GAL4/mal-d^Δ7^* flies. Interestingly, driving the *UAS-MRTFB^R104G^* variant in the *mrtf-T2A-GAL4/mal-d^Δ7^* background proved to be lethal. To further explore the effect of our *MRTFB^R104G^* and *MRTFB^A91P^* variants, we elected to overexpress the *MRTFB^Ref^* and variant UAS lines using a variety of well-established Gal4 drivers (FIG 3A). Our trials using two different ubiquitous drivers (actin- and tubulin-Gal4) demonstrated that broad overexpression of both our *MRTFB^Ref^* and variant lines was lethal (FIG 3B). Expression of either line also failed to demonstrate any morphological differences when expressed in the eyes using GMR-Gal4 (FIG 3C). The expression of *MRTFB^R104G^* within the central nervous system using *Nsyb* and *elav-Gal4* lines did result in partial adult lethality, but the expression of the *MRTFB^A91P^* variant using these drivers produced near-total lethality when crossed with *elav-GAL4* but no reduction in viability when crossed with *Nsyb-GAL4* (FIG 3B). Surviving *MRTFB^R104G^* flies were tested for behavioral deficits using bang-sensitivity assays and climbing assays, but no changes in behavior were identified in either assay (FIG 3D–E). However, when we overexpressed human *MRTFB^Ref^* or either the *MRTFB^R104G^* or *MRTFB^A91P^* variant using a driver expressed in the developing pouch, *nubbin-Gal4*, we observed a significant change in wing morphology (FIG 4A) when the flies were raised at 25°C. In animals overexpressing the human *MRTFB^Ref^* cDNA, the changes were confined to the posterior crossvein, which displayed a highly-penetrant shortening of the vein. Conversely, animals expressing the human *MRTFB^R104G^* or *MRTFB^A91P^* variant displayed a significant disruption of wing morphology, including wing blistering, expansion of intervein tissue, proximal wing vein thickening, shortening of the longitudinal veins, and loss of the anterior and posterior cross veins. Neither of these effects was observed when *nubbin-Gal4* was used to drive the expression of a UAS-LacZ line. Similar effects on wing morphology could also be observed at 18°C and 29°C. At 18°C, animals overexpressing the *MRTFB^Ref^* line displayed a similar foreshortening of the posterior crossvein, but a forking structure could be observed at the terminus of the crossvein. Likewise, the *MRTFB^R104G^* and *MRTFB^A91P^* variants resulted in similar changes to wing morphology at 18 degrees but lacked the wing blistering observed at higher temperatures. At 29°C, flies overexpressing the *MRTFB^Ref^* UAS line displayed identical morphological changes to what was observed at 25°C, while flies expressing the *MRTFB^R104G^* and *MRTFB^A91P^* variants demonstrated further wing damage with extensive blistering, complete loss of wing veins and thickening of the wing margins at the higher temperature (FIG 4A). No changes in wing morphology were observed in animals expressing the *UAS-LacZ* construct at either 18°C or 29°C. As the transcriptional activity of *MRTFB* is highly dependent on its ability to bind its cofactor SRF^27,28^, we wanted to further investigate how co-overexpression of human *MRTFB* and the *Drosophila* ortholog to human *SRF*, *blistered* (*bs*), would effect wing morphology and survival. In *Drosophila, bs* is required for vein and intervein formation within the wings and promotes the development of intervein tissue^5^. Wings taken from *UAS-MRTFB^Ref^*; *nubbin-Gal4 / UAS-bsORF* flies exhibited significant morphological changes, including significant blistering and a near total loss of features within the wing including the loss of all longitudinal veins and crossveins (FIG 4B). By comparison, *UAS-MRTFB^R104G^*; *nubbin-Gal4 / UAS-bsORF* and *UAS-MRTFB^A91P^*; *nubbin-Gal4/ UAS-bsORF*animals did not survive. These findings, taken in aggregate, suggest that the *MRTFB^R104G^* and *MRTFB^A91P^* variants act as either a gain of function or in a dominant negative manner.

**Figure 3.**
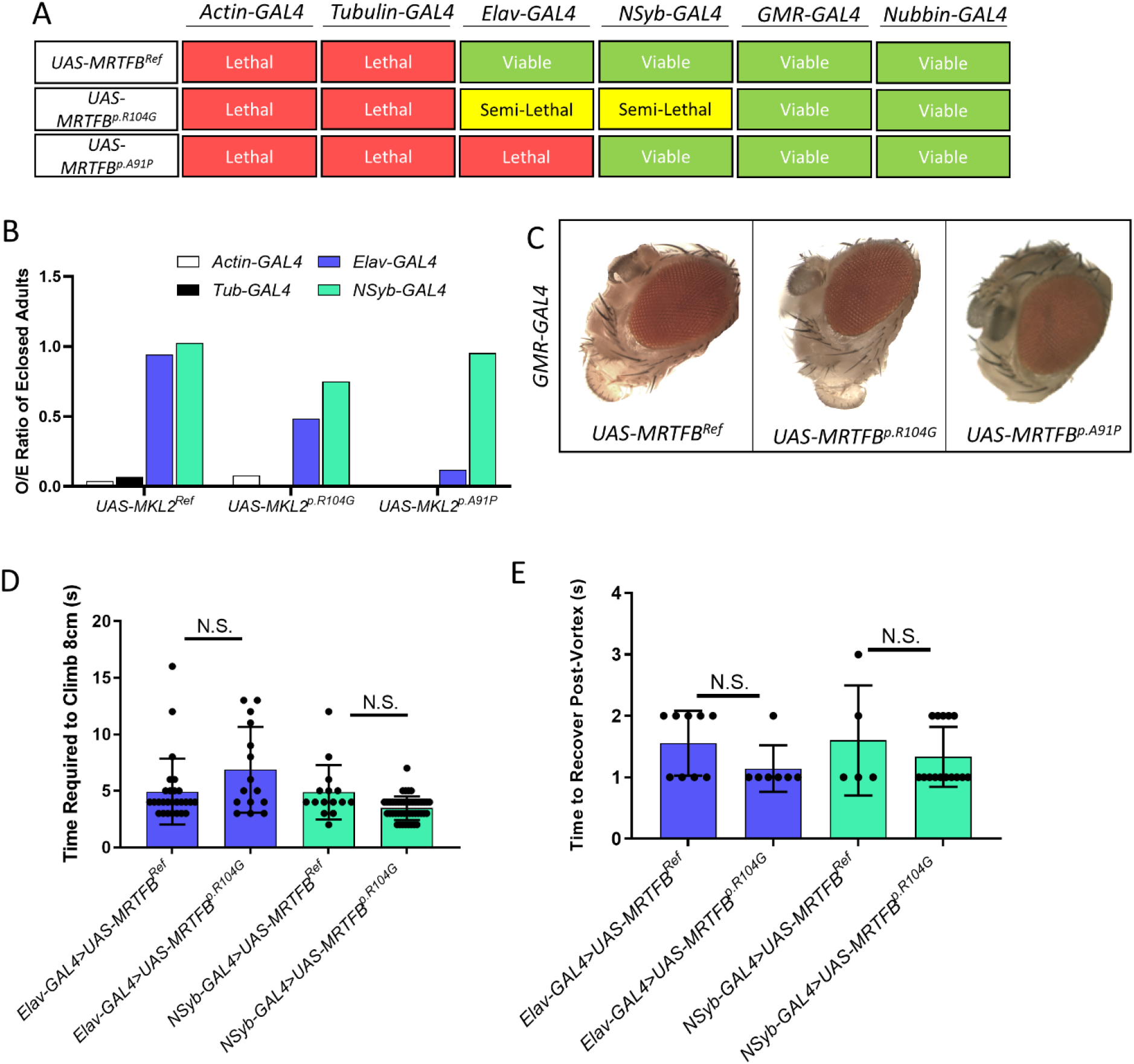
**(A)** Diagram indicating viability when *UAS-MRTFB^Ref^, MRTFB^p.R104G^*, and *MRTFB^p.A91P^*flies are crossed to indicated GAL4 lines. Of note, all crosses to ubiquitous drivers (Actin and Tubulin) are lethal, while neuronal drivers (Elav and Nsyb) display reduced survivability when crossed to *MRTFB^p.R104G^*, and *MRTFB^p.A91P^* lines. **(B)** Ratio of the number of observed flies to the number of expected flies for four different GAL4 drivers. Crosses with *UAS-MRTFB^p.R104G^*, and *UAS-MRTFB^p.A91P^* lines do not generate significant numbers of progeny when paired with actin or tubulin, and *UAS-MRTFB^p.R104G^* flies display semi-lethality when crossed to Elav-GAL4 or Nsyb-GAL4. Crosses between *UAS-MRTFB^rpA91P^* flies and Elav-GAL4 animals display an even greater level of lethality, but these results are not recapitulated in crosses with Nsyb-GAL4 animals. **(C)** Representative images of eyes taken from *GMR-GAL4; UAS-MRTFB^Ref^, GMR-GAL4; MRTFB^p.R104G^*, and *GMR-GAL4; MRTFB^p.A91P^* animals. No significant differences could be identified between reference and variant animals. **(D)** Climbing assays performed at 15 days post-eclosure. No significant differences were observed between *Elav-GAL4; UAS-MRTFB^p.R104G^* and *Elav-GAL4; UAS-MRTFB^Ref^* or between *Nsyb-GAL4; UAS-MRTFB^p.R104G^* and *Nsyb-GAL4; UAS-MRTFB^Ref^* animals. **(E)** Bang sensitivity assays performed at 15 days post-eclosure. No significant differences were observed between *Elav-GAL4; UAS-MRTFB^p.R104G^* and *Elav-GAL4; UAS-MRTFB^Ref^* or between *Nsyb-GAL4; UAS-MRTFB^p.R104G^* and *Nsyb-GAL4; UAS-MRTFB^Ref^* animals.

**Figure 4.**
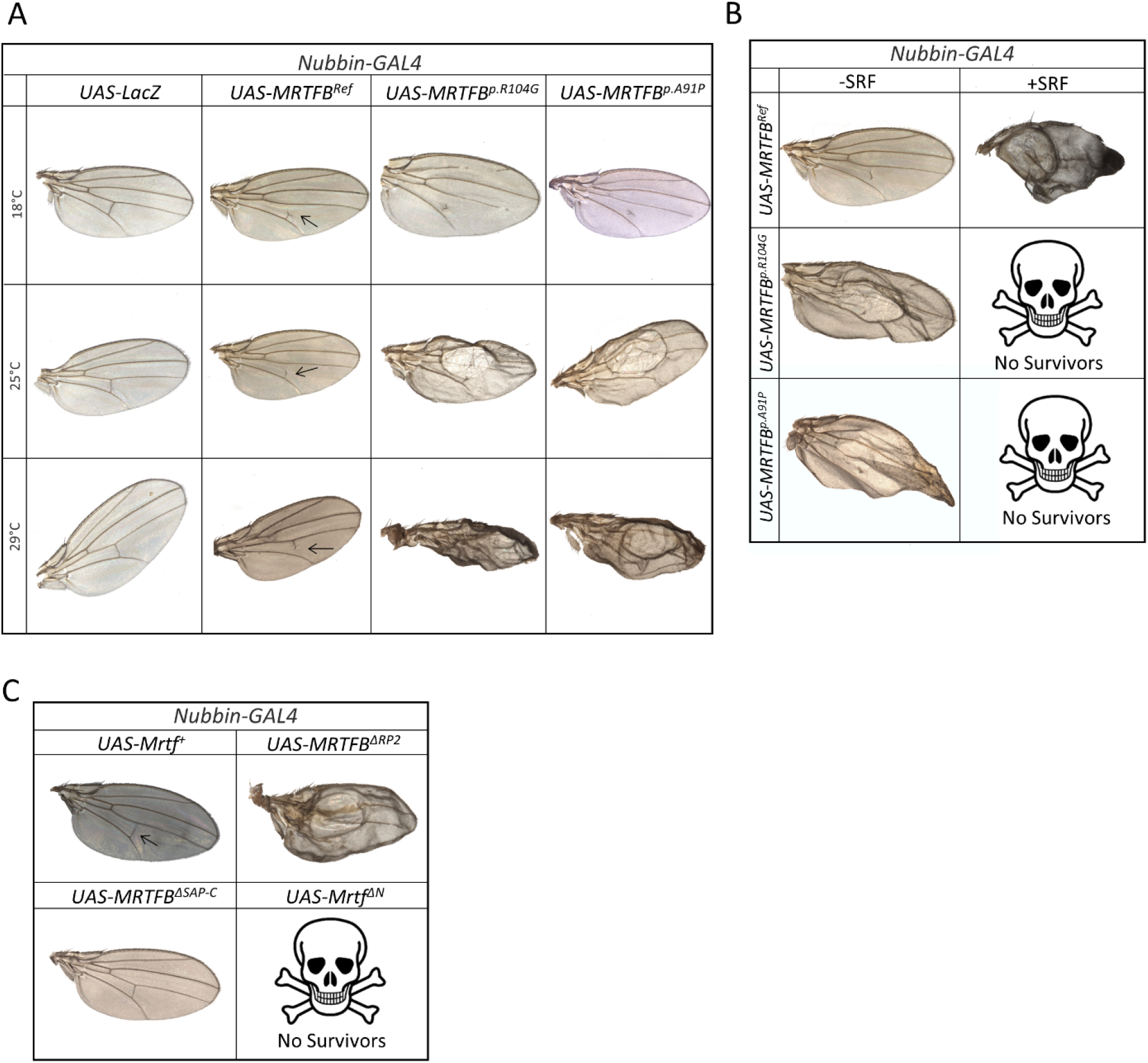
**(A)** Representative images of wings taken from *UAS-MRTFB^Ref^/Nubbin-GAL4, UAS-MRTFB^p.R104G^/Nubbin-GAL4, UAS-MRTFB^p.A91P^/Nubbin-GAL4 and UAS-LacZ/Nubbin-GAL4* animals raised at 18°C, 25°C, and 29°C. *UAS-LacZ* expressing animals showed no significant phenotype at any temperature. Flies expressing *UAS-MRTFB^Ref^* showed truncations in the posterior crossvein. Animals expressing *UAS-MRTFB^p.R104G^* or *UAS-MRTFB^ppA91P^* displayed a significant change in wing morphology which increased in severity as the temperature increased. **(B)** Co-overexpression of MRTFB and its cofactor SRF (*blistered*). The overexpression of *UAS-MRTFB^Ref^* or the *UAS-MRTFB^p.R104G^* or *UAS-MRTFB^p.A91P^* variants alone produced wing damage indistinguishable from what had been previously observed (left column). The coexpression of *UAS-MRTFB^Ref^* and SRF resulted in wings lacking veins and significant blistering. Co-overexpression of SRF and either the *UAS-MRTFB^p.R104G^* or *UAS-MRTFB^p.A91P^* variants was lethal. **(C)** Overexpression of truncated MRTFB using a Nubbin-GAL4 driver. Overexpression of Mrtf^ΔN^ (lacking the first 163 amino acids) causes lethality (left column). The effect of overexpressing fly *mrtf* is indistinguishable from the overexpression of *UAS-MRTFB^Ref^*, causing only minor changes to the posterior crossvein. Likewise, the expression of our Mrtf^ΔC^ line caused no significant changes in morphology in the fly wing, while overexpression of *UAS-MRTFB^δRP^* resulted in wing damage similar to what we have reported for the *UAS-MRTFB^p.R104G^* and *UAS-MRTFB^p.A91P^* variants.

As the overexpression of Drosophila *Mrtf/bs* is known to have an effect on the development of wing tissue^43,44^, we compared the morphological effects caused by our human lines with changes that might occur when Drosophila *Mrtf* is overexpressed in the wing. Using the Nubbin-Gal4 driver to overexpress a *UAS-Mrtf* line resulted in truncations to the posterior cross vein, similar to what was observed when our *MRTFB^Ref^* line was overexpressed. The N-terminal domain of *Mrtf* contains the highly conserved RPEL domains, which are known to be essential for the proper sequestration of the protein in the cytoplasm^15^. A UAS line containing a truncated version of *Mrtf (MrtfΔN)* lacking the first 163 amino acids was overexpressed using the same *nubbin-Gal4* driver as used in previous experiments. Overexpression of the *MrtfΔN* line results in pupal lethality, suggesting that proper regulation of *Mrtf* is essential for survival (FIG 4C). However, as our *MRTFB^R104G^* and *MRTFB^A91P^* variants retain two putatively active RPEL domains, we hypothesized that the ablation of the 2^nd^ RPEL domain alone would result in wing damage but not lethality. To explore this, we generated a *UAS-MRTFB^ΔRP2^* which lacks the 2^nd^ RPEL domain. When the *UAS-MRTFB^ΔRP2^* line was crossed with a *nubbin-Gal4* driver, the resulting progeny displayed significant wing defects, including wing blistering, expansion of intervein tissue, proximal wing vein thickening, shortening of the longitudinal veins, and loss of the anterior and posterior cross veins (FIG 4C). These changes were reminiscent of the alterations in morphology observed in flies expressing the *MRTFB^R104G^* and *MRTFB^A91P^* variants and demonstrate that they are likely disrupting the function of the RPEL domain. However, to rule out the possibility that the observed effects of our variant are the result of a dominant negative mechanism, we generated a truncated version of *MRTFB* lacking the last 422 amino acids (*MRTFB^ΔSAP-C^*). Previously, it has been shown that truncated *Mrtf* lacking its c-terminal region lacks transcriptional activity and has a dominant negative effect. We therefore overexpressed our *MRTFB^ΔSAP-C^* line using Nubbin-Gal4 and observed no differences between animals overexpressing our truncated human protein and animals expressing LacZ (FIG 4C). This suggests that the morphological changes caused by the overexpression of the *MRTFB^R104G^* and *MRTFB^A91P^* variants are not due to a dominant negative effect but are instead due to a change or lack of the function of the 2^nd^ RPEL domain.

As *MRTFBs* activity as a transcriptional coactivator is regulated in part by actin binding within the N-terminal RPEL domains, we hypothesized that the morphological changes driven by the overexpression of the *MRTFB^R104G^* and *MRTFB^A91P^* variants was the result of changes in actin binding to the protein. To evaluate what effect our variant may have, we generated a truncated version of human *MRTFB^Ref^* and *MRTFB^R104G^* bound to a MBP tag. This construct could then be exposed to monomeric G-Actin and assessed for its Actin binding potential. Previous experiments have demonstrated that *MRTFB* displays a high affinity for Actin binding^45,46^. We therefore exposed these truncated constructs, containing the three RPEL domains, to a gradient of Actin concentrations ranging from 1uM to 20uM. As expected, Actin binding to our reference *MRTFB* construct was robust, increasing with the concentration of actin, up to 10μM. Conversely, the *MRTFB^R104G^* variant construct showed significantly reduced levels of actin binding at all concentrations of G-Actin exposure (FIG 5A). To ensure that these findings were also applicable to full-length versions of the protein, we extracted *MRTFB^Ref^, MRTFB^R104G^* and *MRTFB^A91P^* variant protein from flies following heat-shock based induction of expression. The immunoprecipitated proteins were then stained for Actin to determine if bound Actin could be detected. As before, Actin binding could be observed in all three constructs, but the Actin binding ability of *MRTFB^R104G^* and *MRTFB^A91P^* variant protein was significantly reduced (FIG 5B). These results demonstrate that both patient variants have both reduced capacity and ability to bind Actin, which has a direct effect on the regulation of *MRTFB*.

**Figure 5.**
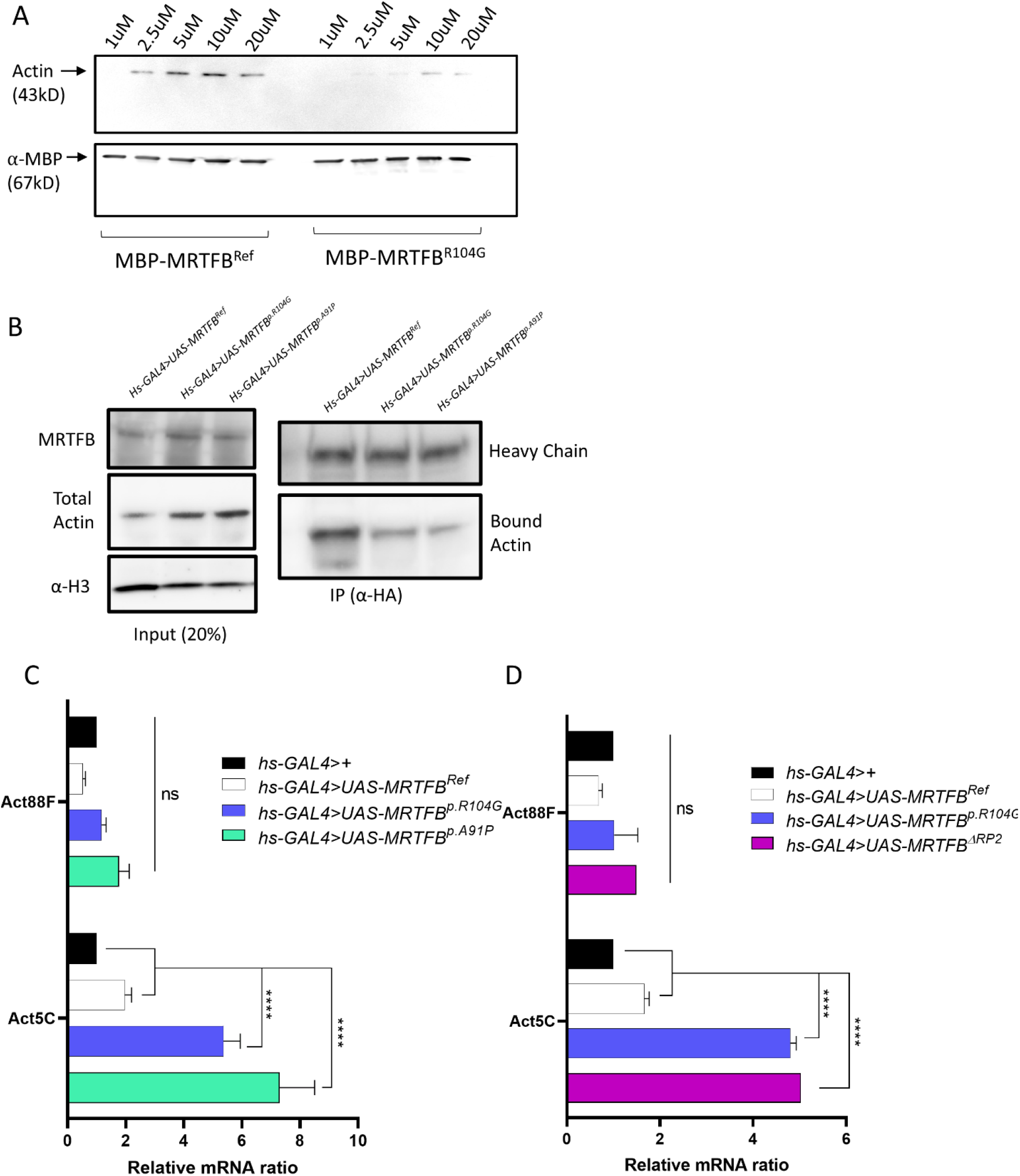
**(A)** Actin binding assay. Truncated human MRTFB^Ref^ and MRTFB^R104G^ bound to a MBP tag was exposed to actin in concentrations ranging from 1uM to 20uM. Actin binding to MRTFB^Ref^ was robust and could be observed at concentrations of 2.5uM and above, while actin binding to the MRTFB^R104G^ variant was significantly reduced and even at high concentrations. α-MBP serves as a loading control. **(B)** Immunoprecipitation of MRTFB^Ref^, MRTFB^R104G^, and MRTFB^p.A91P^ protein from 35 fly heads. Both reference and variant protein was HA tagged. Total fraction indicates a slight increase in total actin levels in MRTFB^R104G^ and MRTFB^p.A91P^ flies. MRTFB^R104G^ and MRTFB^p.A91P^ expressing animals also demonstrate a decrease in the amount of bound actin when immunoprecipitated. **(C)** qPCR measurement of Act88F and Act5C levels post-heat shock in MRTFB^Ref^, MRTFB^R104G^, and MRTFB^p.A91P^ animals. No meaningful change in Act88F was observed. The expression of MRTFB^Ref^ resulted in a 1.5 fold increase in Act5c levels, while the expression of MRTFB^R104G^, and MRTFB^p.A91P^ caused a 5-7 fold increase in Act5c. All measurements were normalized to *hs-GAL4/+* flies. **(D)** qPCR measurement of Act88F and Act5C levels post-heat shock in MRTFB^Ref^, MRTFB^R104G^, and MRTFB^ΔRP2^ animals. No meaningful change in Act88F was observed. The expression of MRTFB^Ref^ resulted in a 1.5-fold increase in Act5c levels, while the expression of MRTFB^R104G^, and MRTFB^ΔRP2^ caused a 5-fold increase in Act5c. All measurements were normalized to *hs-GAL4/+* flies.

When MRTFB is not bound to Actin, it is able to translocate to the nucleus, where in combination with its cofactor SRF contributes to the regulation of an estimated 300 different genes via interaction with a CArG box DNA element^23,47–49^. As our *MRTFB^R104G^* and *MRTFB^A91P^* variants display significantly reduced Actin binding, it is likely to cause alterations in some or all of its downstream targets. In order to verify that the reduced actin binding displayed by our variant was sufficient to have a functional impact on gene expression, we examined the transcription levels of several genes known to be targets of the MRTFB/SRF complex. As *Mrtf* signaling has been shown to be critical for Actin regulation in *Drosophila^27^*, we first examined the effect of overexpressing *MRTFB^Ref^, MRTFB^R104G^*, and *MRTFB^A91P^* on *Actin 5c (Act5c)* transcripts. *Act5c* is orthologous to mammalian ActB, which is a known target for mammalian SRF^22^. The overexpression of *MRTFB^Ref^* resulted in a ~1.5 fold increase in *Act5c* levels when compared to control animals, while *MRTFB^R104G^* and *MRTFB^A91P^* expression increased *Act5c* transcripts by nearly 5 fold (FIG 5C). Overexpression of our *UAS-MRTFB^ΔRP2^* also resulted in a 5-fold increase in *Act5c* expression, similar to what was observed in our *MRTFB* variant animals (FIG 5D). As *Mrtf* has also been implicated in the regulation of a number of different Actin binding proteins, we also wanted to examine the effect of variant *MRTFB* overexpression on a subset of these orthologs. Of the genes chosen, neither the expression of *MRTFB^Ref^* or *MRTFB^R104G^* caused any significant changes in the expression levels of Actin binding proteins, including *twinstar (tsr), chickadee (chic), or Actin-related protein 3 (Arp3)*, nor in the expression of endogenous *Mrtf* or the SRF ortholog *blistered (bs)* (FIG S1 A). These experiments confirm that the expression of *MRTFB^R104G^* and *MRTFB^A91P^* have functional consequences for the regulation of actin itself, and possibly other genes as well, but additional work is required to fully map the transcriptional consequences of the *MRTFB^R104G^* and *MRTFB^A91P^* variants. Furthermore, the changes observed in animals overexpressing *MRTFB^R104G^* or *MRTFB^A91P^* are not significantly different from animals overexpressing *MRTFB^ΔRP2^*, confirming that the change in regulation is dependent on the activity of the 2^nd^ RPEL domain. To further explore the impact of this change on the Actin cytoskeleton, we used the *hs-flp; Act>y+>Gal4, UAS-GFP / SM6A* line to conditionally overexpress *MRTFB^Ref^, MRTFB^R104G^*, and *MRTFB^A91P^* constructs in a subset of *Drosophila* follicle cells. Two days following the initial heat shock, ovaries were dissected and stained with phalloidin, allowing us to directly compare the F-Actin distribution in cells expressing the cDNA with non-expressing cells (FIG 6). We observed that cells expressing either the *UAS-MRTFB^R104G^* or *UAS-MRTFB^A91P^* lines displayed increased F-Actin staining, suggesting that the lack of MRTFB regulation has functional consequences for organization of the Actin cytoskeleton.

**Figure 6.**
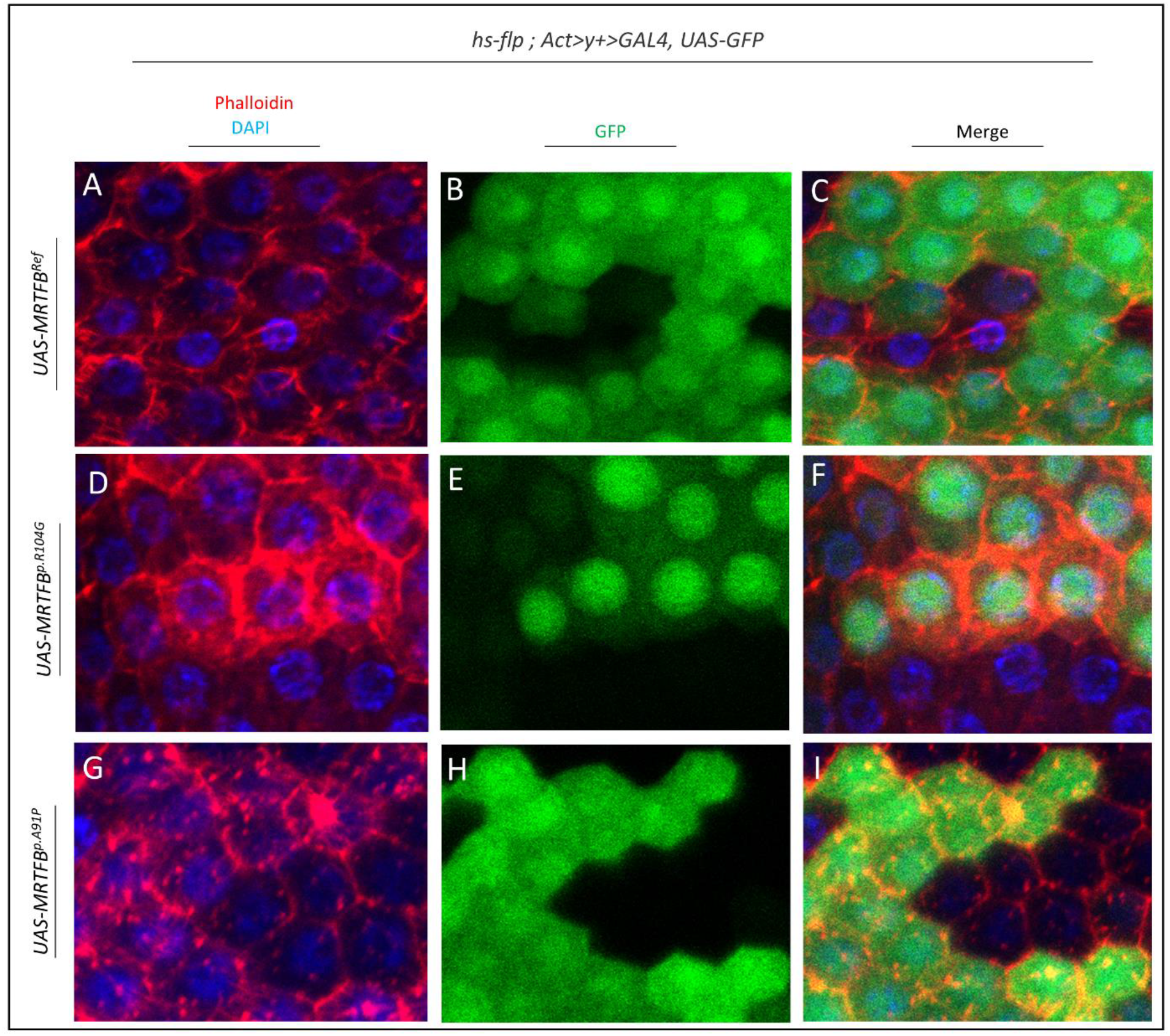
**(A-I)** Representative images take from ovaries of *hs-flp; Act>y+>GAL4, UAS-GFP/UAS-MRTFB^Ref^, hs-flp; Act>y+>GAL4, UAS-GFP/UAS-MRTFB^p.R104G^*, and *hs-flp; Act>y+>GAL4, UAS-GFP/UAS-MRTFB^p.A91P^*flies following one-hour heat shock. **(A, D, G)** Phalloidin and DAPI staining of ovaries. Actin overgrowth can be clearly seen in *hs-flp; Act>y+>GAL4, UAS-GFP/UAS-MRTFB^p.R104G^*, and *hs-flp; Act>y+>GAL4, UAS-GFP/UAS-MRTFB^p.A91P^* flies. **(B, E, H)** GFP indicating regions producing GAL4 post heat shock. **(C, F, I)** Merge demonstrating that the highest levels of actin staining can be observed in regions producing GAL4.

Beyond the two probands reported here, we made an effort to identify additional *MRTFB* variants, and requested variants from Baylor genetics. We found two variants (*MRTFB^N95S^* and *MRTFB^E109Q^*) within the RPEL domain in cases that had not been previously solved. However, these variants were not *de novo*, both being inherited from phenotypically normal parents. As with our *MRTFB^Ref^, MRTFB^R104G^* and *MRTFB^A91P^* lines, we elected to overexpress the *MRTFB^N95S^*and *MRTFB^E109Q^* lines using the same battery of Gal4 drivers as used prior to evaluate any changes in morphology that may result from overexpression. Similar to what we had observed previously, overexpression of any of the lines with an *Actin* or *tubulin-Gal4* driver was lethal at 25°C, and no changes to eye morphology was observed as the result of expression using a *GMR-Gal4* driver (FIG S2 A, C). Likewise, neuronal overexpression using a *Nsyb-* or *elav-Gal4* line did not result in any significant changes in viability (FIG S2 B). Using *nubbin-Gal4* to drive the expression of either the *MRTFB^N95S^* and *MRTFB^E109Q^* variant lines resulted in a truncation in the posterior cross vein, similar to what was observed in *UAS-MRTFB^Ref^*; *nubbin-Gal4* or *UAS-mrtf; nubbin-Gal4* animals (FIG S2 D). In both *MRTFB^N95S^*and *MRTFB^E109Q^* expressing animals, ~60% of flies displayed a forked end at the truncation of the posterior crossvein when raised at 18*°*C or 25°C, while flies raised at 29°C did not display a forked truncation. Therefore, the *MRTFB^N95S^* and *MRTFB^E109Q^* variants do not exhibit the functional consequences seen in the *MRTFB^R104G^* and *MRTFB^A91P^* cases. These findings suggest that the *MRTFB^R104G^* and *MRTFB^A91P^* variants are specifically disruptive to the function of *MRTFB*, and that the location of the variant within the RPEL domain is extremely important when considering its potential to disrupt the function of the protein. While variants outside of the RPEL domains may also have functional consequences to the functionality of the *MRTFB* protein, their effects are yet to be determined in our model system.

## Discussion

Here we describe two individuals with *de novo* variants in *MRTFB* which result in a neurodevelopmental phenotype that includes intellectual disability, speech apraxia, impulse control issues, gross and fine motor impairments, and dysmorphic facial features including low-set posteriorly rotated ears, a depressed nasal bridge, epicanthal folds, midface hypoplasia, and down-slanting palpebral fissures. Prior to this study, *MRTFB* had not been categorized as a human disease gene but had been studied in the context of both fatal human microcephaly and cancer. In the single case study associated with microcephaly, a speculative model was established based on three children from the same parents with a nonsynonymous deleterious variant in the basic domain of *MRTFB* and a 185kb deletion in cis with the variant allele^31^. The authors acknowledge that their results did not causally demonstrate the link between fatal microcephaly and *MRTFB* and additional cases have not as yet been identified. Here, we describe individuals with overlapping phenotypes and demonstrate that the variants in *MRTFB* reported here lead to dramatic differences in *MRTFB* regulation. Our findings represent clinical and functional aspects within a specific domain of *MRTFB*. It is possible that alterations in other domains or deletions could contribute to other pathologies. The studies implicating *MRTFB* as an oncogene also reveal that *MRTFA/MRTFB* was found to promote epithelial-mesenchymal transition and to contribute to cell motility through regulation of cell adhesion and cell spreading in human cell lines^50,51^. Examination of human tumors has also revealed an upregulation of *MRTFA/MRTFB*, as well as a *RREB1-MRTFB* and *Cllorf95-MRTFB* fusion genes in chondroid lipomas and ectomesenchymal chondromyxoid tumors respectively^52–54^. While our current results pertain to germline, not somatic variants in *MRTFB*, further studies will be needed to explore whether the lack of regulation by Actin may affect whether germline variants alter metastatic processes.

Prior mouse models of *MRTFB* also display different defects depending on the allele used for the study. LacZ enhancer trap alleles have been shown to result in vascular defects and perinatal lethality in mouse models, while deletions of *MRTFB* resulted in complete lethality between E13.5 and E14.5^6,25,26^. The cause of mortality in these cases has been attributed to abnormalities in vascular development, with distinct defects present in the cardiac outflow tract and branchial arch arteries. These defects have been attributed to a lack of differentiation in smooth muscle cells. One of the consequences of this altered state of vascular development is liver defects, an abnormal extension of the liver parenchyma, and hemorrhage during development in mouse models. Interestingly, *MRTFB’s* cofactor *SRF*, is also known to produce liver expansion in constitutively active *SRF* mouse models^55^. As proband 1 was diagnosed with hepatomegaly, it is possible that the *MRTFB^R104G^* gain-of-function variant identified here may result in heightened levels of *SRF* activity leading to this phenotype, although further research is necessary to confirm this.

While our study indicates that human *MRTFB* may not be fully able to replace the fly ortholog *Mrtf*, functional differences can be observed in our model between our reference and variant lines when overexpressed using a *nubbin-Gal4* driver. As lethality can be observed in *MRTFB^R104G^* and *MRTFB^A91P^* animals during our humanization experiments, and co-expression of *blistered*, the fly ortholog of *SRF*, resulted in increased lethality in our variant, *MRTFB^R104G^* and *MRTFB^A91P^* both appear to act as gain of function alleles. While we cannot fully rule out the possibility that our variants represent neomorphic alleles, a gain of function model where increased *MRTFB/SRF* activity is driven by a lack of actin regulation in our variants is consistent with both the current state of knowledge concerning *MRTFB* as well as our own experiments (FIG 7). The gain of function nature of the allele appears to be highly position dependent within the RPEL domain, as the *MRTFB^N95S^*and *MRTFB^E109Q^* variants tested within the domain did not reveal any significant changes within the wing tissue. Our Actin binding assay also demonstrates that the *MRTFB^R104G^* and *MRTFB^A91P^* variants have a significant effect on the ability of *MRTFB* to bind Actin, although it is currently unclear if this is due to a lack of Actin binding within the second RPEL domain or a change in the larger pentameric Actin-MRTFB complex. This lack of regulation by Actin appears to be the key to several of the phenotypes observed in the probands, possibly including the neurological phenotypes. As evidenced by mouse and rat models, proper *MRTFB/SRF* activity is important for the production of dendritic spines, establishing proper dendritic complexity, and assembling neural circuits^4^. While additional research is necessary to demonstrate if and how *MRTFB^R104G^* and *MRTFB^A91P^* affect neurological development, it is plausible that the lack of regulation present in these alleles may be sufficient to alter normal patterns of neuronal growth and development.

**Figure 7.**
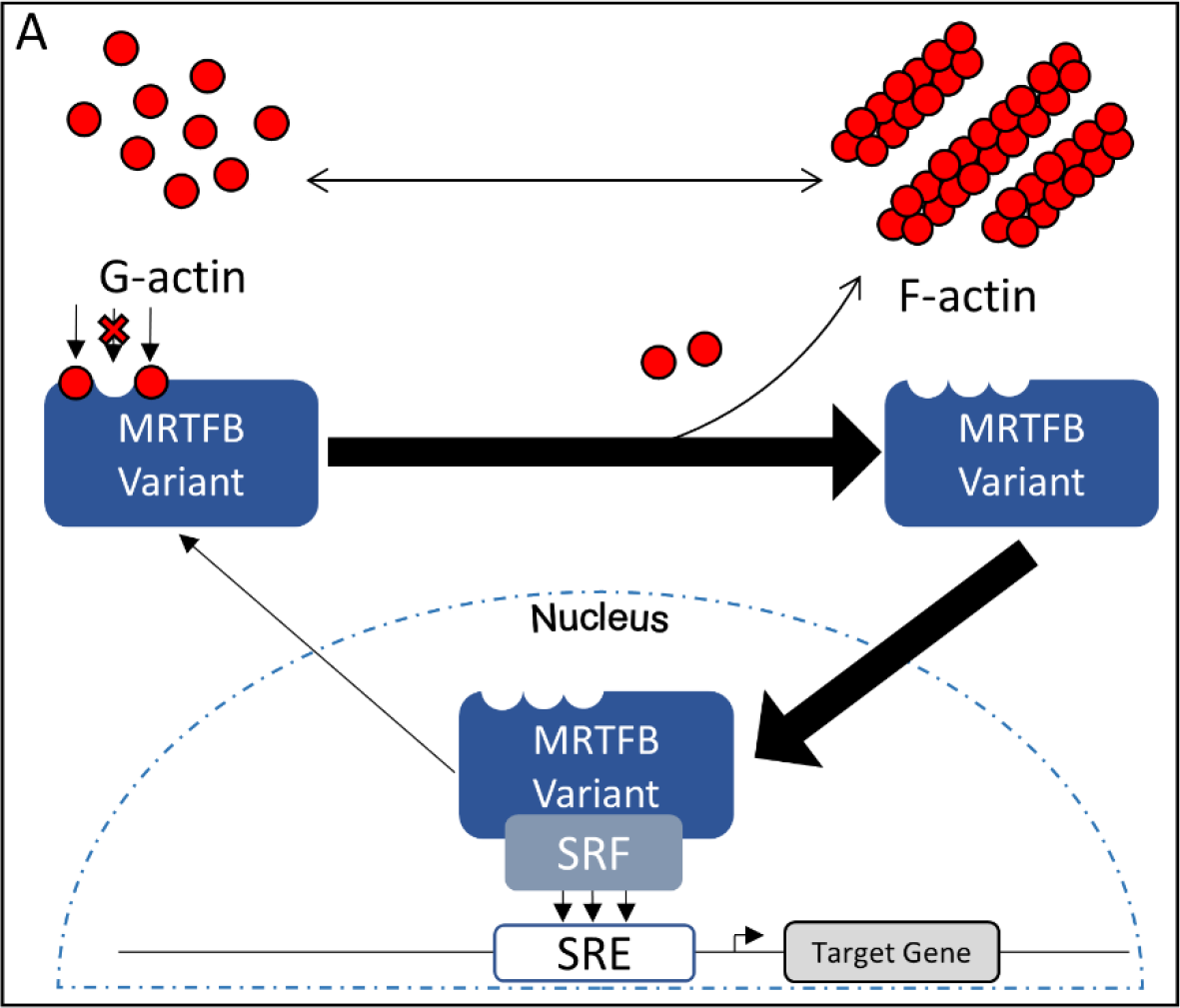
(A) Proposed model of MRTFB-related disorder. The presence of a variant within the 2^nd^ RPEL domain results in a reduction in MRTFB/actin binding. This change in the regulation of MRTFB results in a higher rate of nuclear translocation and activity with its cofactor, SRF. This in turn results in a higher rate of target gene transcription, including transcription of *Actin5c*.

In summary, we have identified two probands with significant phenotypic overlap and *de novo* missense variants within the 2nd RPEL domain of *MRTFB*. Using a *Drosophila* model based upon these variants, we established that expression of *MRTFB^R104G^* or *MRTFB^A91P^* using a *Mrtf-T2A* resulted in lethality, whereas expression of the *MRTFB^Ref^* control did not. Likewise, expression of *MRTFB^R104G^* or *MRTFB^A91P^* within wing tissues resulted in significant damage to the fly’s wing. Our results indicate that the *MRTFB^R104G^* and *MRTFB^A91P^* variants likely have a gain of function mechanism. These changes can also be observed when *MRTFB^ΔRP2^* was overexpressed, indicating that the *MRTFB^R104G^* and *MRTFB^A91P^* variants effectively disrupted the RPEL domain preventing it from interacting with Actin. This observation was further validated using an Actin binding assay which demonstrated that the *MRTFB^R104G^* allele was ineffective at binding actin at all concentrations tested. This inability to regulate transcriptional activity through Actin binding had a significant effect on Act5c levels, with flies expressing either *MRTFB^R104G^* or *MRTFB^ΔRP2^* displaying a 5-fold increase in Actin level. These findings highlight a possible means of pathogenesis, specifically that a lack of Actin binding results in a loss of Actin-based regulation leading to a change in transcriptional activity. In the future, follow-up studies examining the effect of the *MRTFB^R104G^* and *MRTFB^A91P^* allele on transcriptional targets will help us to better understand what other genes are misregulated in our probands, and further exploration using human neuronal cell lines or *Drosophila* brain tissue may shed light on the neurological effects observed in our proband.

## Accession Numbers

The variants in this study have been submitted to ClinVar, with the following accession numbers: NM_014048.4; c.310C>G; p.R104G corresponds to SCV002599424. NM_014048.4; c.270G>C; p.A91P corresponds to SCV002599423.

## Acknowledgments

This paper is supported by NIH grants U54 NS 093793 (The UDN MOSC) to M.W and S.Y., 1F32 NS110174-01 Fellowship award to J.C.A, and NRF-Korea Fellowship NRF-2021R1A6A3A14044510 awarded to J-W.M. We are thankful for the resources provided by the Bloomington *Drosophila* Stock Center and the Developmental Studies Hybridoma Bank for their contribution of fly stocks and antibodies.

## Materials and Methods

### Patient recruitment and consent

Probands were recruited by their referring physicians at local sites (NIH and UCLA). Proband 1 (UDN061684) was identified through the Undiagnosed Disease Network (UDN), while Proband 2 was integrated into the study through genematcher/UDN. Prior to research, informed written consent was obtained for testing and publication according to the standards and practices of the institutional review boards and ethics committees of each institution. Documents and forms used in the consent process were standardized according to the standards of the UDN.

### *Drosophila* Stocks

| **Stock** | **Source** |
| --- | --- |
| *UAS-CD8::GFP* | **Bellen Laboratory** |
| *y[1] w[\*]; P{w[+mC]=Act5C-GAL4}17bFO1/TM6B, Tb[1]* | **BDSC #3954** |
| *y[1] w[\*]; P{w[+mC]=tubP-GAL4}LL7/TM3, Sb[1] Ser[1]* | **BDSC #5138** |
| *w; GMR-GAL4* | **Bellen Laboratory** |
| *hs-flp ; Act>y+>Gal4, UAS-GFP / SM6A* | **Bellen Laboratory** |
| *w[1118]; M{w[+mC]=nSyb-GAL4.P}ZH-86Fb/TM6B* | **BDSC #68222** |
| *P{w[+mC]=GAL4-elav.L}2/CyO* | **BDSC #8765** |
| *Nubbin-Gal4* | **Bellen Laboratory** |
| *Mrtf[Delta7] P{ry[+t7.2]=neoFRT}80B/TM6B, Tb[1]* | **BDSC #58418** |
| *w[\*]; Tl{w[+mW.hs]=Tl}Mrtf[ko] P{ry[+t7.2]=neoFRT}80B/TM6B, Tb[1]* | **BDSC #58419** |
| *Mrtf-T2A-Gal4/TM3 Sb* | **This study** |
| *y w; UAS- MRTFB (Reference)/SM6a* | **This study** |
| *y w; UAS- MRTFB (p.R104G)/SM6a* | **This study** |
| *y w; UAS-MRTFB (p.N95S)/SM6a* | **This study** |
| *y w; UAS-MRTFB (p.E109Q)/SM6a* | **This study** |
| *y w; hs-Gal4* | **Gift from K-W Choi** |

### *Drosophila* housing and handling

All flies were grown in a temperature and humidity-controlled incubator at either 18, 25 or 29 degrees Celsius as indicated in the text. All animals were maintained at 50% humidity on a 12-hour light/dark cycle. Flies were allowed to feed on a standard fly food medium consisting of water, yeast, soy flour, cormeal, agar, corn syrup and propionic acid. Collection of animals for experiments was performed during daylight hours.

### Generation of *Mrtf-T2A-Gal4* line

CRISPR-mediated insertion of the T2A-Gal4 cassettes was accomplished as previously described (Kanca et al, 2019, Kanca et al, 2022). Briefly, a *Mrtf* specific plasmid containing a restriction cassette flanked on either side by a 200-nucleotide homology arm and a gRNA1 target sequence, gRNA1 coding sequence, and gene specific sgRNA (targeting the site TCAATCGAGGGTGTTGGATCTGG) was synthesized (Genewiz) in a pUC57_Kan_gw_OK2 vector backbone. A swappable integration cassette containing *attP-FRT-T2A-GAL4-polyA-3XP3-EGFP-FRT-attP* was subsequently subcloned into the plasmid, replacing the restriction cassette. 250 ng/μl of the completed plasmid was then injected into *y w; iso; attP2(y+){nos-Cas9(v+)}* embryos. The resulting G0 males and females were crossed to *y w* flies to screen for the presence of *3XP3-EGFP*. Two independent lines were successfully established and PCR validated through single fly PCR (Gloor et al., 1993) using OneTaq PCR master mix (NEB #M0271L). Primers for PCR were designed to flank the integration site (primer sequences: *Mrtf_*forward ACTCAGAGCAACGTTGATCACG, *Mrtf*_reverse: ATTGTCTCAGACGACGATGGCA) and were used in combination with insert-specific primers (primer sequences: CRIMIC_ch_rev: GCGGAAGAGAGATAAATCGGTTG and CRIMIC_ch_for: GTGGTATGGCTGATTATGATCAGAAG) that bind 5’ of the cassette in reverse orientation and 3’ of the cassette in forward orientation.

### Generation of UAS-human cDNA lines

UAS lines were generated as previously described^56^. Briefly, reference human cDNA clones corresponding with NM_014048.4 were obtained in pcDNA3.1+/C-(K)DYK vector through genescript. Clone was amplified through PCR via MangoMix (Meridian Bioscience) and cloned into a pDONR221 destination vector using BP clonase enzyme. Variants were generated in the pDONR vector via site-directed mutagenesis utilizing NEBaseChanger with the Q5 mutagenesis kit (NEB) and verified by sequencing. LR clonase II (ThermoFisher) enzyme was then used to shuttle sequence verified reference and variant ORFs into the p.UASg-HA.attB destination vector. Verified ORF’s in the p.UASg-HA.attB vector were microinjected into ~200 embryos and transferred into a attP docking site (VK00037) via ΦC31 mediated transgenesis.

### Immunofluorescence staining and confocal microscopy

Adult brains were submerged in 4% paraformaldehyde (Electron Microscopy Sciences) in PBTX for 30 minutes post dissection. Fixed brains were washed 3 times to remove residual paraformaldehyde then incubated overnight at 4°C in donkey normal serum (DNS), 0.2% PBTX, and primary antibodies on a shaker. After 24 hours, brains were again washed 3 times, and again incubated overnight at 4°C in DNS, 0.2% PBTX and secondary antibodies on a shaker. Brains were then washed a final 3 times in PBS and mounted in Vectashield mounting medium (Vector Labs, H-1000-10) for imaging. Images were obtained with a laser confocal microscope (Zeiss LSM 880) with 20X objective, and image processing was accomplished using ZEN (Zeiss) and ImageJ software. Primary antibodies used: Mouse anti-repo (DSHB: 8D12) 1:200 and Rat anti-elav (DSHB: 7E8A10) 1:200. Secondary antibodies used: Anti-mouse 647 (Jackson ImmnoResearch 715-605-151) 1:200, Anti-rate Cy3 (Jackson ImmunoResearch 712-165-153) 1:200.

### Imaging of adult morphology

Imagining of whole adults expressing GFP was performed under CO2 anesthesia using a Leica DMC6200 with pE-300 CoolLED and LasX software. *mal-d^Δ7^ mutant crosses were also evaluated using the* Leica DMC6200 after flies were frozen at −20°C for at least 24 hours. *Drosophila* wing and heads were removed under CO2 anesthesia and placed onto a slide for imaging. Images were obtained Leica MZ16 stereomicroscope equipped with an Optronics MicroFire Camera and Image Pro Plus 7.0 software to perform extended-depth-of-field images.

### Bang sensitivity and climbing assays

Flies to be used were isolated 1-3 days post-eclosure and group housed for 10 days until assessment. On the day of the trial, flies were transferred into an empty polystyrene vial via aspiration. Climbing assays were performed by tapping the flies to the bottom of the vial 3 times by hand, then observing the flies as they climbed towards a marker on the vial 8 cm above the vials base. The time required to reach the marker was recorded via digital stopwatch. Any flies which failed to reach the marker within 60 seconds were recorded as requiring 60 seconds, and the trial stopped. Immediately after the climbing assay, a bang sensitivity assay was performed. In the same vial, flies were vortexed at full speed (Fisher STD Vortex Mixer, Cat. No, 02215365) for 10 seconds and recovery times recorded using a digital stopwatch. Both bang and climbing assays were performed on a minimum of 20 flies.

### Assessment of lethality and morphological phenotypes

Crosses for assessment of lethality and morphological phenotypes were performed using-Gal4 drivers as indicated in the text, using 5-10 virgin females crossed to a similar number of males. Parents were transferred to a new vial after 5-7 days so as collect multiple generations of progeny. After 3 passages, any surviving parents were discarded. Flies were collected after most pupae eclosed, and the total number of flies scored based on the presence or absence of balancers. For lethality assessment, a minimum of 70 flies were scored. Viability was calculated via evaluation of the number of observed progeny compared to the number of expected progeny based on mendelian ratio. Animals were classified as lethal if the O/E ratio was less than .15, and semi-lethality is classified as an O/E ratio less than .8. Assessment of morphological phenotypes was only done for animals lacking balancers, and phenotypes were noted if they appeared in more than 70% of progeny.

### MBP fusion protein expression and purification

For MBP fusion protein purification, the expression vector (pMal-c5x) was transformed into the competent cell [XJb (DE3) autolysis competent cells, T3051, Zymo research]. After overnight incubation (37 °C, LA media, 200 rpm), cells were inoculated (1:100) into fresh LA media with sterilized L-arabinose (final concentration: 3mM). Add IPTG (final concentration: 0.2mM) for protein induction when OD value reaches 0.6~0.8. After 2.5 hours of incubation at 37 *°C* with shaking (200 rpm), cells were transferred to a 25 *°C* shaking incubator (200 rpm) for 16 hours. Then, cells were harvested at 6,000 g, 4°C for 30 minutes. The supernatant was discarded, and the remaining cell pellet was stored at −20 *°C* fridge for 24 hours to induce the autolysis. Frozen cells were incubated on ice for 15 minutes, then incubated at 25 *°C* for 15 minutes on an orbital shaker. After autolysis, MBP column buffer [200mM NaCl, 20mM Tris-HCl pH 7.4, 1mM EDTA, 1X protease inhibitor cocktail (APExBIO)] was used for pellet resuspension. Cells were centrifuged at 10,000 g 4 *°C* for 10 min, and only the supernatant part was used for further application. Pre-cleared MBP resin (NEB, E8021S) was added to the supernatant and incubated for 16 hours at 4 *°C* at the orbital rotator. After incubation, the resin was harvested by gentle centrifuge (2000 g, 4 °C). The resin was thoroughly washed at least three times (5 min for each washing) with ice-cold MBP column buffer. Then, elution buffer (MBP column buffer + 10mM maltose) was added to the resin.

Resin was further incubated for 5 min at room temperature with constant rotation by the rotator. Eluted proteins were loaded on the centrifugal filter unit (Millipore-Sigma, UFC8030) and centrifuged at 3,500 g 4°C for 20 minutes to eliminate any remaining maltose from the elute. Concentrated proteins were diluted by MBP column buffer (without maltose) and then further centrifuged by filter unit. This elimination step was repeated three times to get maltose-free proteins. Protein purity was determined by Coomassie staining, and proteins with high purity (>90%) were used for the actin-binding assay.

### Actin binding assay

Purified (>99%) human actin monomers (Cytoskeleton, APLC99) were prepared (5 mM Tris-HCl pH 8.0, 0.2 mM CaCl2, 0.2 mM ATP, 5% sucrose, and 1% dextran). MBP fusion proteins (150mM), general actin buffer (5mM Tris-HCl pH 8.0 and 0.2mM CaCl2, Cytoskeleton) were incubated with actin protein (1mM, 2.5mM, 5mM, 10mM, and 20mM respectively) at room temperature with constant rolling. BSA (150mM) was added for negative control instead of MBP fusion protein. After rolling, pre-cleared MBP resin (NEB, E8021S) was added to each tube and further incubated for 16 hours at 4°C. A protease inhibitor cocktail (APExBIO) was added to prevent protein degradation. After incubation, the resin was washed twice with ice-cold MBP column buffer to wash out any unbound proteins. Then, elution buffer (MBP column buffer + 10mM maltose) was added to the resin and incubated at room temperature for 10 min with constant rolling. Eluted proteins were analyzed further by western blot.

### Immunoprecipitation

Flies were raised at room temperature before eclosion. After eclosion, adult flies were given heat-shock at 37°C for 30 minutes. Then, flies were placed at 29°C incubator for 24 hours. Flies were once gain given 30 minutes of heat-shock at 37°C then were lysed with lysis buffer [50mM Tris (pH8.0), 1% NP-40, 1mM EGTA, 150mM NaCl, Protease inhibitor cocktail (APExBIO, K1007)] after 4 hours of further incubation at 29°C incubator. 15 males and 15 female flies were used for the lysis. To perform the lysis evenly, glass homogenizer was utilized. Samples were centrifuged for 10 minutes at 14,000g 4°C. Only supernatant part was collected for future applications. After aliquoting input samples, magnetic bead (Genscript, L00273) was added to the remaining sample for the preclearing (30 minutes at room temperature). After removing beads from the samples, anti-HA magnetic beads (Pierce, 88836) were added to the samples and incubated for 15 minutes at room temperature on rotator. Then, samples were further incubated overnight at 4°C. After the incubation, beads were washed with 0.1% NP-40 in PBS for at least three times using magnetic rack (Invitrogen, 12321D). Beads were treated with Laemmli Sample buffer (Bio-Rad, 1610737) to extract bound proteins.

### Western Blot

Samples taken from prior assays were mixed 1:1 with Laemmli Sample Buffer (Bio-Rad) with 10% 2-mercaptoethanol and vortexed to ensure adequate mixing. The resulting mixture was centrifuged for 30 seconds, then heated on an aluminum block at 90°C for 5 minutes. Heated samples were allowed to cool, then centrifuged for 10 minutes at 14,000 RPM. Samples were then loaded into a premade 4-20% gradient gel (bio-rad). A PVDF (polyvinylidene difluoride) membrane was activated via the application of 100% methanol for 10 seconds. After running, the gel was transferred to PVDF via wet transfer, and the PVDF membrane was blocked using a 5% blocking solution (1x TBST with 0.1% TWEEN-20 and 2.5g nonfat dry milk or BSA) for 1 hour at room temperature with rotation. The membrane was then incubated overnight at 4°C with anti-H3 [1:5,000 (07-690, Sigma-Millipore)], anti-actin [1:2,000 (MAB1501, Sigma-Millipore)], anti-MBP [1:2000 (66003-1-Ig, Proteintech)], or MKL2 Rabbit pAb anit-MKL2 (1:5000, ABclonal) respectively. The membrane was washed at least three times with 0.1%TBST before the 2^nd^ antibody incubation. SuperSignal™ West Pico PLUS (34580, Thermo) and Western Lightning Plus-ECL (NEL104001EA, Perkin Elmer) were used as chemiluminescent substrates. Detection was performed using a Bio-Rad ChemiDoc MP imaging system.

### QPCR

Adult *hs-Gal4/+; UAS-MRTFB (Reference)/+, hs-Gal4/+; UAS-MRTFB (p.R104G)/+*, and *hs-Gal/+; +/+* flies were isolated and stored at 29°C. One day prior to mRNA extraction, the flies were exposed to a 37°C heat shock for a period of 30 minutes, then returned to their incubator. The day of the experiment, flies were again exposed to 37°C heat shock for a period of 30 minutes, returned to the incubator for a period of 4 hours before extraction. 10 flies were selected, and total RNA was extracted via TRIzol (Fischer) following the manufacturer’s recommendation. cDNA synthesis was subsequently performed using iScript (Bio-Rad) as per manufacturer instructions. Normalized concentrations of cDNA obtained through reverse transcription were then used for quantitative PCR experiments using a Bio-Rad CFX96 Real-Time System and iQ SYBR Green Supermix (Bio-Rad). Primers with a high primer efficiency (>95%) were chosen for amplification, with sequence information as follows: 1) MRTFB Fwd-CTCCTGTCCTCCCCACAAAC, Rev-GTCCATCTGCGGCTCATTCT; 2) Act5C Fwd-AAGTACCCCATTGAGCACGG, Rev-ACATACATGGCGGGTGTGTT; 3) Act88F Fwd-TCGATCATGAAGTGCGACGT, Rev-ACCGATCCAGACGGAGTACT; 4) *mrtf* Fwd-CAGACAGTCACCACCAAAGG, Rev-GTCGCACCATGAGCTTCACTT; 5) *tsr* Fwd-ATGGCTTCTGGTGTAACTGTG, Rev-TGACATAGCGATGCTTTTTGTCC; 6) *chic* Fwd-ATGAGCTGGCAAGATTATGTGG, Rev-TCCTCTTTTGTCACCTCAAAGC; 7) *bs* Fwd-TACACGACCTTCTCCAAGCG, Rev-GTTGAGGCAGGTCTGGATGA, 8) *rp49* Fwd-TACAGGCCCAAGATCGTGAA, Rev-TCTCCTTGCGCTTCTTGGA; 9) *arp3* Fwd-ATTTGCCGGGAATAAAGAGCC, Rev-CGCGCAGATACTTAAAGACGC. All primers were generated using PrimerBlast or were verified primers validated by the DRSC Primerbank. All primers used for qPCR were purified via high-performance liquid chromatography, and were optimized to function at a Tm of 60°C.

### Ovary Dissections

*hs-flp; Act>y+>GAL4, UAS-GFP/UAS-MRTFB^Ref^, hs-flp; Act>y+>GAL4, UAS-GFP/UAS-MRTFB^p.R104G^*, and *hs-flp; Act>y+>GAL4, UAS-GFP/UAS-MRTFB^p.A91P^* flies were heat shocked at 37°C for 10 minutes 24 hours prior to ovary dissection. Adult ovaries were dissected in PBS, fixed in 4% paraformaldehyde for 20 minutes, and then rinsed three times in 2% PBTx. Fixed ovaries were incubated with Alexa anti-phalloidin-568 antibody (diluted 1:300) for 15 minutes, washed three times in 2% PBTx and mounted in Vectashield with DAPI (Vector Labs, H-1200). Images were obtained with a laser confocal microscope (Zeiss 710) with 20X objective, and image processing was accomplished using ZEN (Zeiss) software.

### Web Resources

ClinVar: https://www.ncbi.nlm.nih.gov/clinvar/

CADD: https://cadd.gs.washington.edu/

DIOPT: https://www.flyrnai.org/cgi-bin/DRSC_orthologs.pl

ExAC: http://exac.broadinstitute.org/

Genematcher: https://genematcher.org/

Geno2MP: https://geno2mp.gs.washington.edu/Geno2MP/#/

gnomAD: https://gnomad.broadinstitute.org

MARRVEL: http://marrvel.org

OMIM: https://omim.org/

PolyPhen-2: http://genetics.bwh.harvard.edu/pph2/

