## Supplemental Figure for "*De Novo* Variants in *MRTFB* have gain of function activity in *Drosophila* and are associated with a novel neurodevelopmental phenotype with dysmorphic features"

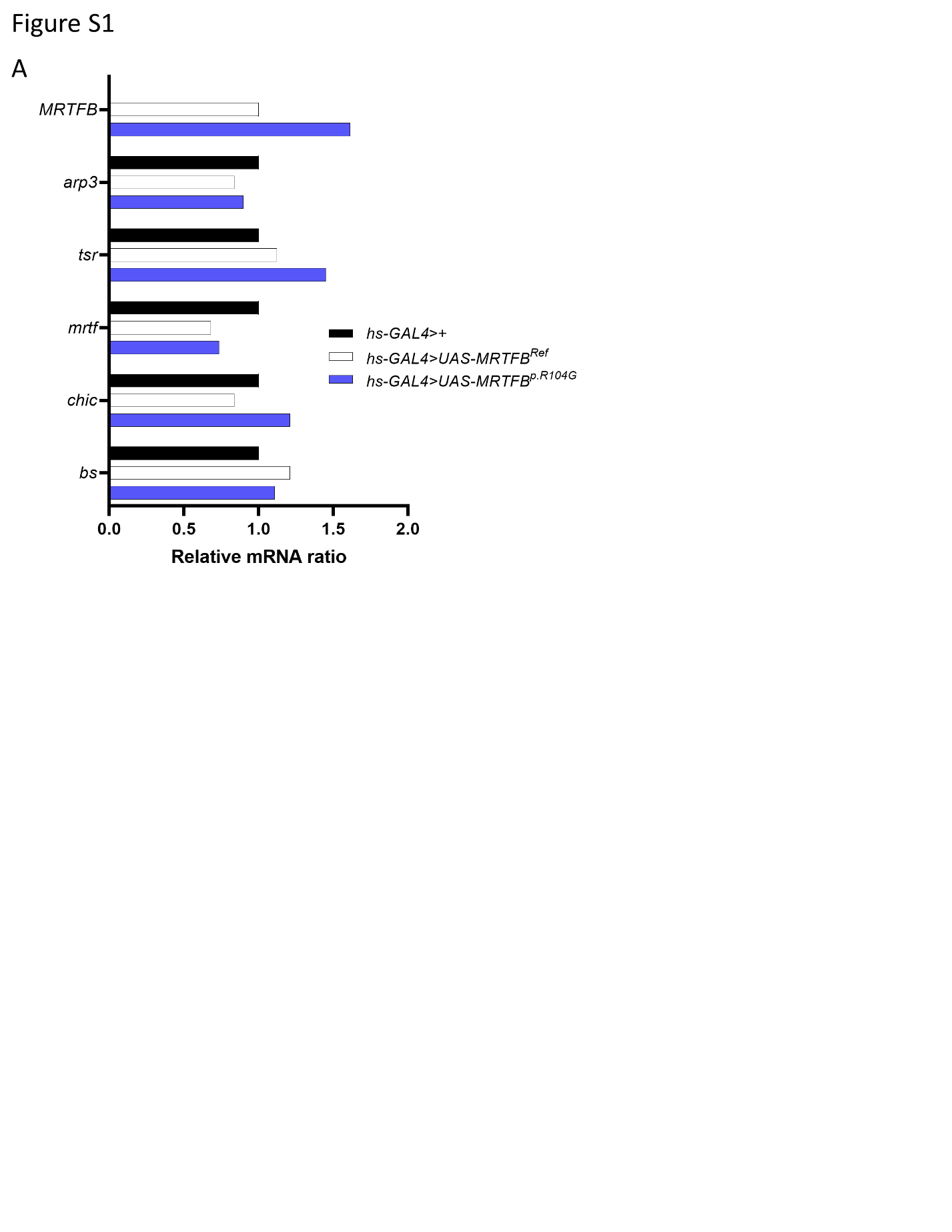

Figure S1

**(A)** qPCR measurement of actin-associated proten levels post-heat shock in *Hs-GAL4*/+, *Hs-GAL4; UAS-MRTFB^Ref^*, and *Hs-GAL4 ; UAS-MRTFB^R104G^* animals. No meaningful change in the levels of *mrtf, chic, tsr, arp3, chic,* or *bs* was observed. Human MRTFB could only be observed when driven by the *hs-GAL4* line.

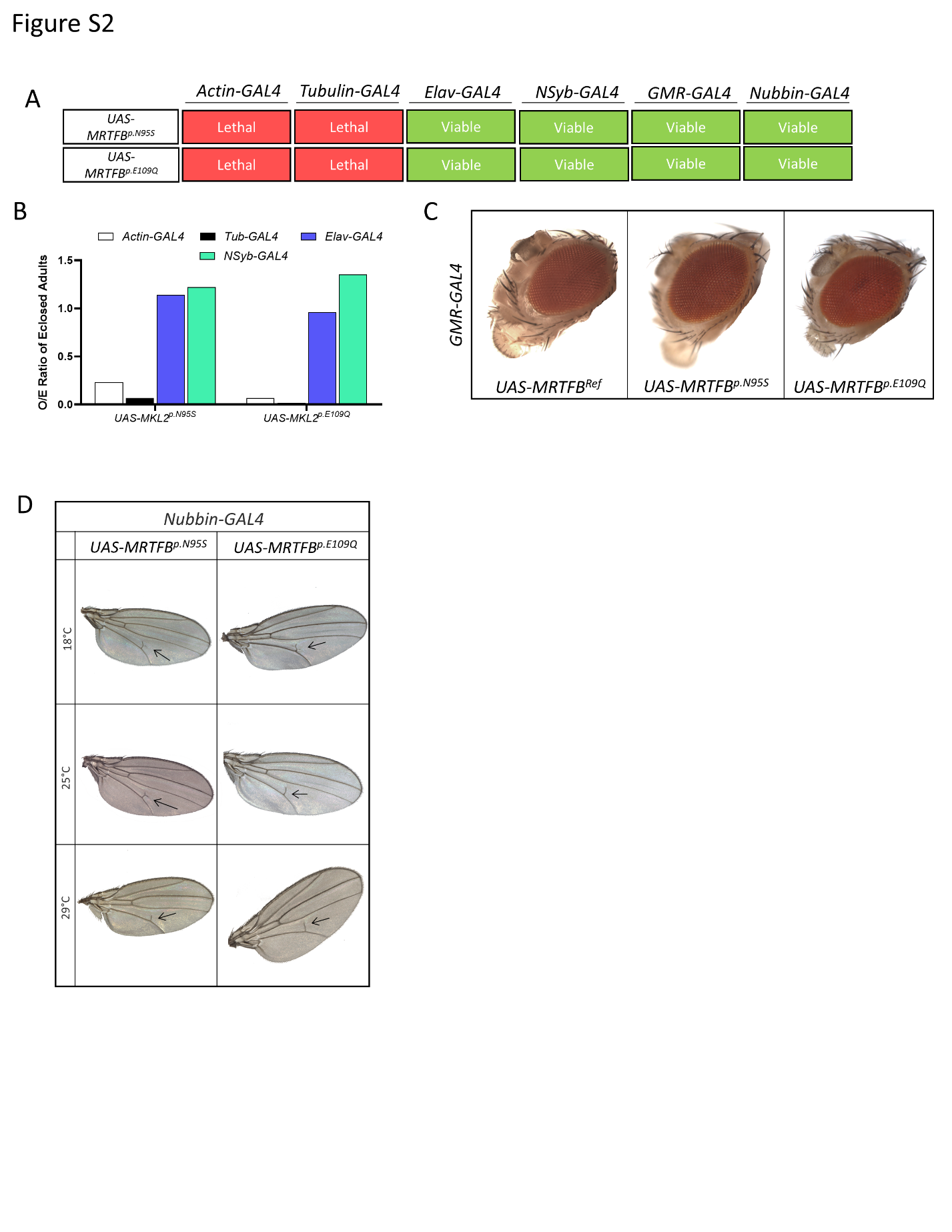

Figure S2

**A)** Diagram indicating viability when MRTFB^p.N95S^, and MRTFB^p.E109Q^ flies are crossed to indicated GAL4 lines. Of note, all crosses to ubiquitous drivers (Actin and Tubulin) are lethal, while all other drivers produce viable offspring. **(B)** Ratio of the number of observed flies to the number of expected flies for four different GAL4 drivers. Crosses with MRTFB^p.N95S^ and MRTFB^p.E109Q^ demonstrate expected levels of progeny when crossed to neuronal drivers, and crosses against actin or tubulin do not generate significant numbers of progeny. **(C)** Representative images of eyes taken from *GMR-GAL4 ; UAS-MRTFB^Ref^*, *GMR-GAL4 ;* MRTFB^p.N95S^, and *GMR-GAL4 ;* MRTFB^p.E109Q^ animals. No significant differences could be identified between reference and variant animals. **(D)** Representative images of wings taken from *UAS- MRTFB^p.N95S^/Nubbin-GAL4*, *UAS-MRTFB^p.E109Q^ /Nubbin-GAL4* animals raised at 18°C, 25°C, and 29°C. Wings showed slight truncations

Supplemental Text 1

**Proband 1 (FIG. 1B)** is a 12-year-old female who was born at 39 weeks gestation to a healthy 33-year-old G2P1 mother and 41-year-old father of Greek ancestry. Her newborn exam was unremarkable and she was discharged at 2 days of life. Crawling occurred at 12 months, walking at 18 months, and first words at 30 months. At 3 years of age a neurology evaluation diagnosed language delay, gross and fine motor delays, hypotonia and dysmorphic facial features (Table 1). Chromosomal microarray performed at this time revealed a maternally inherited duplication at 20p.12.1.

On examination at the NIH at the age of 12-years and 9-months, she was found to have dysmorphic features including mild synophrys, slightly low-set ears, widely-spaced teeth and inverted nipples, bilaterally. At this time she was also noted to have coarse hair with premature graying (Table 2). Abdominal ultrasound also identified an enlarged liver, although hepatic function tests were normal. Neuropsychological testing provided a diagnosis of intellectual disability, in the moderate-to-severe range, as well as apraxia and severely delayed expressive language. Audiologic testing showed normal peripheral hearing bilaterally, and ophthalmological evaluation was normal save for mild anisometropia/astigmatism. Her neurologic exam revealed brisk reflexes without clonus but was otherwise unremarkable. Notably, she has no history of seizures or regression. At evaluation, she had a normal brain MRI and MRS, although her EEG showed bilateral independent temporal epileptiform discharges, present only during slow wave sleep, suggesting an increased risk for seizures. She has displayed a history of difficult sleep with multiple nighttime awakenings.

Additional genetic evaluation included lysosomal enzyme screening, quantitative mucopolysaccharides, Sanfilippo enzyme testing, and carbohydrate deficient transferrin testing, all of which were normal.

**Proband 2 (FIG. 1C)** is an 11-year-old male with autism spectrum disorder, ADHD, anxiety, cognitive impairments, characteristic facial features, strabismus, and hypotonia (Table 1). He was born at 39 weeks via C-section to a 35- year-old G2P0 mother and 39-year-old father. He was delivered via C-section due to fetal distress and had a 9-day NICU stay for meconium aspiration without requiring intubation. Birth weight of 3180 grams (Z score -0.3), length of 50 cm (Z score +0.1), and head circumference of 34 cm (Z score -0.4). His growth parameters continue to be between the 50th and 75th percentile. He had recurrent otitis media requiring tympanostomy tubes at 14 months. He had surgical repair of his strabismus at age 2 and a febrile seizure at age 3. At age 9, he had a prolonged hospital stay with severe pneumonia with empyema and pleural effusion requiring chest tube placement.

He was noted to have developmental delays at 6 months with rolling at 9 months, walking at 20 months, and first words at 3.5 years. He has had intensive developmental therapies from 9 months of age. A diagnosis of autism spectrum disorder was given at 3 years old. At age 11, he is potty trained, makes good eye contact with adults, struggles with motor planning and can't follow the rules of a game, often has uncontrollable giggling, tantrums, and displays repetitive questioning. He has a history of watching spinning objects and self-stimulatory behaviors with his hands that have resolved. There is no awareness of safety, resulting in an elopement risk. He is athletic and can throw a ball well but struggles with his balance and fine motor skills. He is unable to write or draw and mostly eats with his hands, though he is able to use a fork. He has speech apraxia but communicates verbally in basic sentences. He can read some words but is unable to do math calculations. There have been no developmental regressions.

On physical examination, he has mild synophrys, epicanthal folds, downslanting palpebral fissures, midface hypoplasia, depressed nasal bridge, tubular nose with bulbous tip, low set posteriorly rotated ears, fifth finger clinodactyly, and overlapping toes (Table 2). Other evaluations have included normal CK level and CT abdomen and chest without congenital anomalies. Genetic testing included a normal microarray, very long chain fatty acids, and non-diagnostic mitochondrial genome sequencing. Exome trio sequencing identified a *de novo* variant in MRTFB.
